# Detection of human disease conditions by single-cell morpho-rheological phenotyping of whole blood

**DOI:** 10.1101/145078

**Authors:** Nicole Toepfner, Christoph Herold, Oliver Otto, Philipp Rosendahl, Angela Jacobi, Martin Kräter, Julia Stächele, Leonhard Menschner, Maik Herbig, Laura Ciuffreda, Lisa Ranford-Cartwright, Michal Grzybek, Ünal Coskun, Elisabeth Reithuber, Geneviève Garriss, Peter Mellroth, Birgitta Henriques-Normark, Nicola Tregay, Meinolf Suttorp, Martin Bornhäuser, Edwin R. Chilvers, Reinhard Berner, Jochen Guck

## Abstract

Blood is arguably the most important bodily fluid and its analysis provides crucial health status information. A first routine measure to narrow down diagnosis in clinical practice is the differential blood count, determining the frequency of all major blood cells. What is lacking to advance initial blood diagnostics is an unbiased and quick functional assessment of blood that can narrow down the diagnosis and generate specific hypotheses. To address this need, we introduce the continuous, cell-by-cell morpho-rheological (MORE) analysis of whole blood, without labeling, enrichment or separation, at rates of 1,000 cells/sec. In a drop of blood we can identify all major blood cells and characterize their pathological changes in several disease conditions *in vitro* and in patient samples. This approach takes previous results of mechanical studies on specifically isolated blood cells to the level of application directly in whole blood and adds a functional dimension to conventional blood analysis.

## Introduction

Blood is responsible for the distribution of oxygen and nutrients, and centrally involved in the immune response. Consequently, its analysis yields crucial information about the health status of patients. The complete blood count, the analysis of presence and frequency of all major blood cells, constitutes a basic, routine measure in clinical practice. It is often accompanied by analysis of blood biochemistry and molecular markers reflecting the current focus on molecular considerations in biology and biomedicine.

An orthogonal approach could be seen in the study of the overall rheological properties of blood. It is evident that the flow of blood throughout the body will be determined by its physical properties in the vasculature, and their alterations could cause or reflect pathological conditions (1–3). In this context, blood is a poly-disperse suspension of colloids with different deformability and the flow properties of such non-Newtonian fluids have been the center of study in hydrodynamics and colloidal physics (4). Probably due to the dominant importance of erythrocytes for blood rheology, at the expense of sensitivity to leukocyte properties, this approach has not resulted in wide-spread diagnostic application, maybe with the exception of blood sedimentation rate (5).

Focusing on the physical properties of individual blood cells has suggested a third possibility to glean maximum diagnostic information from blood. Various cell mechanics measurement techniques, such as atomic force microscopy (6–8), micropipette aspiration (1, 9–11) or optical traps (12–14), have been used to show that leukocyte activation, leukemia, and malaria infection, amongst many other physiological and pathological changes, lead to readily quantifiable mechanical alterations of the major blood cells (6, 12, 15–19). These proof-of-concept studies have so far been done on few tens of specifically isolated cells. This line of research has not progressed towards clinical application for lack of an appropriate measurement technique that can assess single-cell properties of sufficient number directly in whole blood.

This report aims to close this gap by presenting a novel approach for high-throughput single-cell morpho-rheological (MORE) characterization of all major blood cells in continuous flow. Mimicking capillary flow, we analyse human blood without any labeling or separation at rates of 1,000 cells/sec. We show that we can sensitively detect MORE changes of erythrocytes in spherocytosis and malaria infection, of leukocytes in viral and bacterial infection, and of malignant transformed cells in myeloid and lymphatic leukemias. The ready availability of quantitative morphological parameters such as cell shape, size, aggregation, and brightness, as well as rheological information of each blood cell type with excellent statistics might not only inform further investigation of blood as a complex fluid. It also connects many previous reports of mechanical changes of specifically isolated cells to a measurement now done directly in whole blood. As such, it adds a new functional dimension to conventional blood analysis — a MORE complete blood count — and, thus, opens the door to a new era of exploration in investigating and diagnosing hematological and systemic disorders

## Results

### Establishment of MORE analysis

In order to establish the normal MORE phenotype of cells found in whole blood, we obtained venous, citrate-anticoagulated blood of healthy donors, of which 50 μl was diluted in 950 μl of measurement buffer with a controlled elevated viscosity, but without any additional labeling, sorting, or enrichment. The cell suspension was then pumped through a micro-channel not unlike micro-capillaries in the blood vasculature (Fig. 1A). Brightfield images of the cells, deformed by hydrodynamic shear stresses in the channel (20), were obtained continuously by RT-DC (21) (see Methods; Movie S1). These images revealed distinct differences in overall morphology, brightness, and amount of deformation between all major cell types found in blood (Fig. 1B). RT-DC further enabled the continuous, real-time quantification of the cross-sectional area and of the deformed shape (see detailed description in Methods and Fig. S1) of an, in principle, unlimited number of cells at measurement rates of 100 – 1,000 cells/sec (Fig. 1C). For each cell detected and analyzed, an image was saved and the average pixel brightness within the cell determined (Fig. 1D, Fig. S1). This single-cell MORE analysis of whole blood revealed distinct and well-separated cell populations in the space spanned by the three parameters (Movie S2). Notably, size and brightness alone — parameters not unlike those accessible by light scattering analysis in standard flow cytometers — were sufficient for the identification of the cell types (Fig. 1D), so that deformation as additional, independent parameter was available for assessing their functional changes. The identity of the individual cell populations by size and brightness was established by magnetic cell sorting, controlled by fluorescence immunophenotyping, and subsequent MORE analysis (Fig.S2). A key feature is the very clear separation of the abundant erythrocytes (red blood cells; RBCs) from other cells as a result of their much greater deformation and lower brightness. This feature gives access to leukocyte properties directly in whole blood, without the potentially detrimental effects of hemolysis or other separation steps, which are required for analysis with cell mechanics techniques with lower specificity and throughput, or non-continuous measurement.

**Fig. 1.**
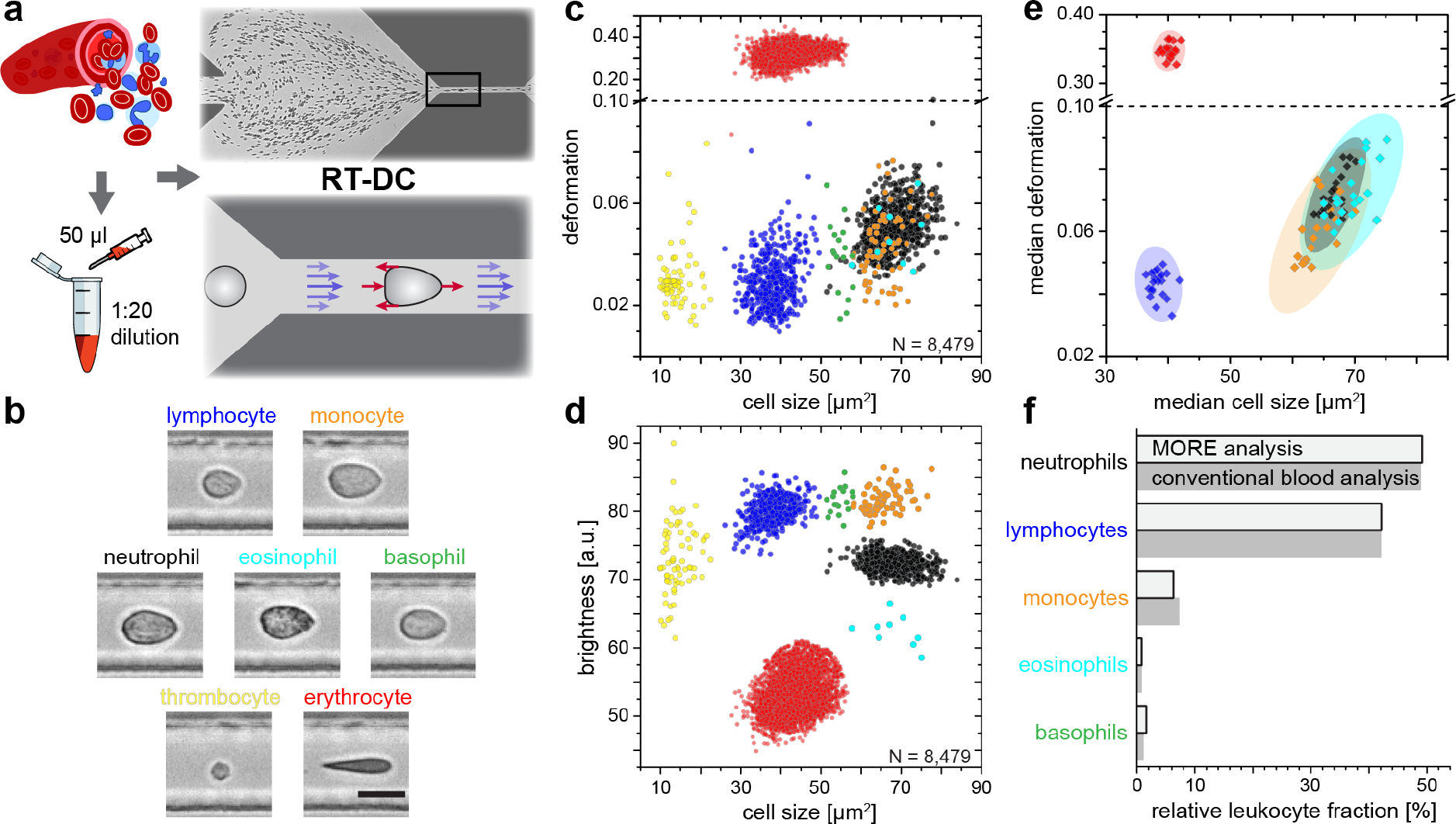
Single-cell, morpho-rheological phenotyping of whole blood. **A**, Analysis of whole, diluted blood. Hydrodynamic shear forces (red arrows) induce deformation of cells passing a microfluidic channel (20 × 20 μm^2^) at speeds of more than 30 cm/s (blue arrows). **B**, Representative images of blood cell types acquired. Scale bar is 10 μm. Images are analyzed for cell size as well as **C**, cell deformation and **D** average cell brightness. Each dot represents one of *N* measurement events. **E**, Normal range of deformation and size of cell populations from healthy donors. Each diamond represents the median of one donor; transparent ellipses indicate 95 % confidence areas. **F**, Comparison of MORE cell counts with conventional blood count.

In extensive tests of the variability of this approach, MORE phenotyping yielded identical results in repeated measurements of blood from the same donor, with sodium citrate added as an anti-coagulant and for different storage times (Fig. S3), between different donors of both sexes (Fig. S4), and blood samples taken at different times during the day (Fig.S5). This robustness served to establish a norm for the different cell types (Fig. 1E). MORE analysis provided the identity and frequency of all major white blood cells as with a conventional differential blood count (Fig. 1F; Table S1) — obtained from a single drop of blood, with minimal preparation, and within 15 min. Going beyond this current gold-standard of routine blood cell analysis, and importantly also beyond all other single-blood-cell mechanical analysis studies to date, MORE phenotyping allowed the sensitive characterization of pathophysiological changes of individual cells directly in whole blood. In the following, we exemplarily demonstrate, in turn for each of the blood cell types, the new possibilities of gaining MORE information from an initial blood test as a time-critical step in generating specific hypotheses and steering further investigation enabled by this approach.

### MORE analysis of erythrocytes

Spherocytosis is a prototypical hereditary disease in humans in which genetic changes (here ankyrin and spectrin mutations) cause abnormal shape and mechanical properties of erythrocytes. Current diagnosis based on shape detection in a blood smear, osmotic fragility assessed by Acidified Glycerol Lysis Time (AGLT) or by osmotic gradient ektacytometry, flow-cytometric determination of staining with Eosin-5-Maleimide (EMA test), or direct detection of the mutation by PCR takes time, requires specific preparation, is costly and does not lend itself to screening. MORE analysis of whole blood of patients with spherocytosis directly revealed significantly stiffer and smaller erythrocytes than normal (Fig. 2A-C). The differences are so clear that this analysis can serve as a fast primary and cheap screening test for spherocytosis to be followed up by more elaborate analysis.

**Fig. 2.**
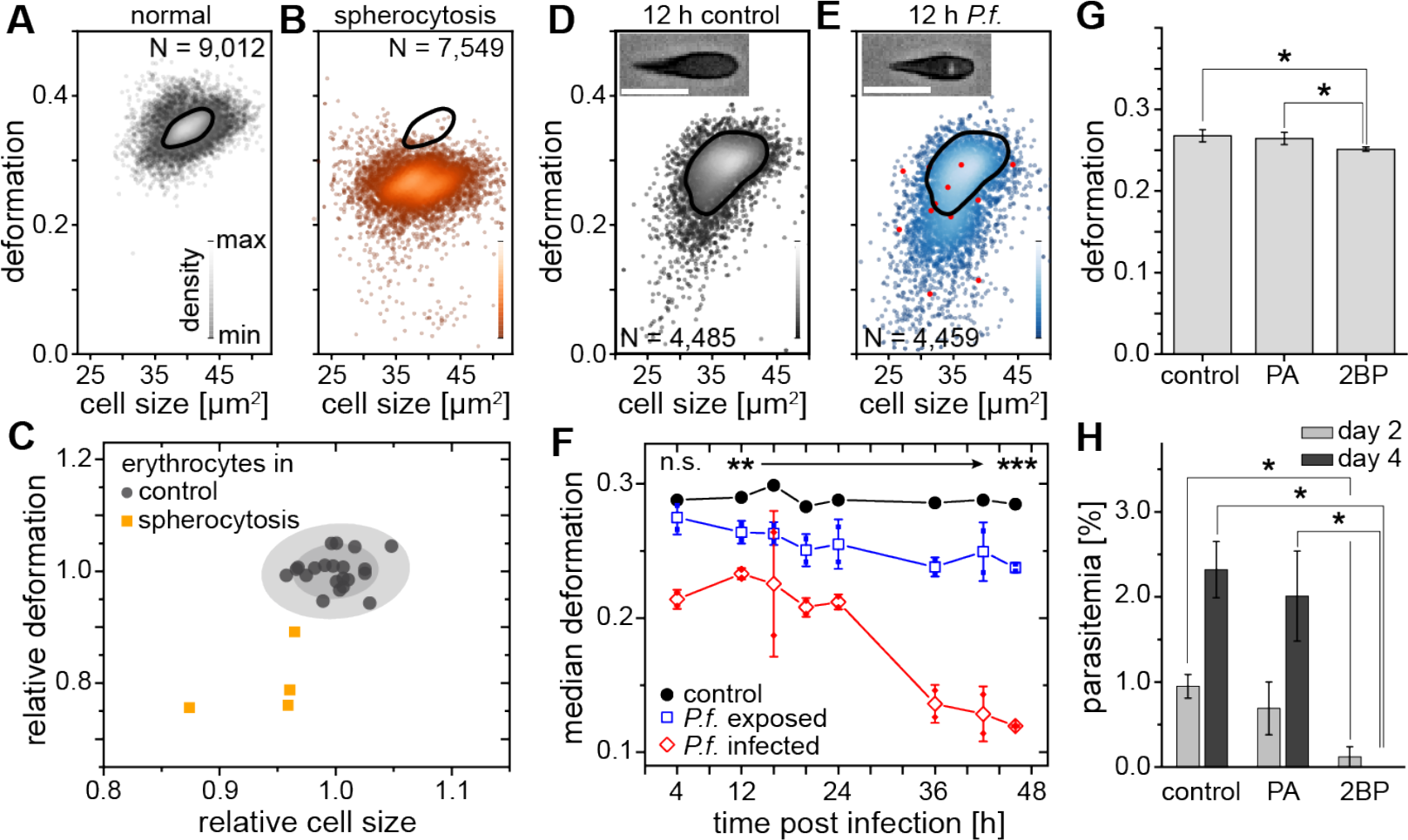
MORE phenotyping detects RBC pathologies. Exemplary density plots of RBC size vs. deformation in samples from **A**, healthy donor and **B**, patient with spherocytosis. **C**, Relative median RBC deformation and size in patients with spherocytosis (orange, *n* = 4) compared to controls (black, *n* = 21 as in Fig. 1E with 68 % and 95 % confidence ellipses). Density plots of size vs. deformation of **D**, control RBCs and **E**, RBCs exposed to *P.f.* (blue, infected RBCs red), both after 12 h incubation. Scale bars, 10 μm. **F**, Evolution of RBC deformation over 46 h time course of control (black), *P.f.* exposed (blue) and *P.f.* infected RBCs (red); open squares and diamonds, mean ± SD, *n* = 2; filled squares, individual medians, ** *p* < 0.01, *** *p* < 0.001. **G**, 2BP-treated RBCs compared to PA- and non-treated controls (mean ± SD of population medians, *n* = 4, * *p* < 0.05). **H**, Reduced parasitemia in 2BP- compared to PA- and non-treated controls at 2 and 4 days post infection. Error bars: SD binomial, * *p* < 0.0125.

A change in RBC deformability has also been implicated in malaria pathogenesis, since single cells infected by parasites have been reported to be stiffer (18). This insight has not progressed towards clinical application and the gold standard in malaria diagnosis is still a manual thick blood smear analysis. To evaluate whether MORE analysis could provide a sensitive, automated alternative, we analyzed populations of RBCs infected *in vitro* with *Plasmodium falciparum* (*P.f.*) with a parasitemia of 7 – 8 % at time points over the 2 day parasite life cycle. We found a clear, significant, and increasing reduction in the deformation of the entire exposed RBC population detectable after 4 h (Fig. 2D-F; Fig. S6). Inspection of the individual cell images revealed the appearance of characteristic features likely associated with the maturation of parasite inside a subset of RBCs (Fig. 2D, E insets; Fig. S6). These features permitted the direct identification of positively infected cells, whose relative frequency peaked at 36 h (Fig. S6). The separate assessment of overtly infected cells showed an even greater deformation reduction than observed in the entire exposed population (Fig. 2F; Fig. S6), which relates to the possibility of clearance of stiff, infected cells from the circulation by the spleen (22, 23). However, this small fraction of stiffer cells alone cannot account for the reduced deformation of the whole population, so that a bystander stiffening of non-infected cells seems involved (14). Inhibition of palmitoylation of membrane proteins by 2-bromo-palmitate (2BP) led to less deformation than in controls and in RBCs treated with palmitic acid (PA; Fig. 2G), with a concurrent reduction in *P.f.* infectivity (Fig. 2H). While a previous report found no change in infectability of RBCs treated with 2BP (24), the difference could stem from the different RBC receptors involved in invasion by the different parasite clones (3D7 vs. HB3), which in turn are differentially affected by palmitoylation. Thus, MORE analysis has the potential to not only simplify, automate, and speed up malaria diagnosis, but also to provide additional quantitative information aiding research into the pathogenesis of the disease (25).

### MORE analysis of leukocytes

While RBC mechanics has already been used for clinical diagnostics using rheoscopes and ektacytometers for over 40 years (15, 22, 26), leukocyte mechanics has not been utilized for diagnostic purposes. This is likely due to their increased stiffness compared to RBCs and a lack of convenient techniques capable of sufficiently deforming them in suspension — their physiological state. Until recently, techniques with sufficient throughput, obviating the need for specifically isolating the relevant cells of interest, which always bears the potential of inadvertent cell change, did not exist. In this sense, the mechanical phenotyping of diagnostic changes of leukocytes directly in whole blood is the most transformative application area of MORE analysis. For example, there have been proof-of-concept studies on the mechanical changes associated with neutrophil activation, with older reports showing a stiffening, in line with the pronounced actin cortex that is a hallmark of neutrophil activation (6, 17). These studies are seemingly in conflict with recent findings reporting a softening of neutrophils with activation (19). While some of the discrepancy could stem from accidental activation by the cell preparation required, the different modes of mechanical testing (11) or the different time-scales of mechanical response assessed by the different techniques, MORE analysis of the neutrophil activation kinetics in whole blood with the bacterial wall-derived tripeptide fMLP suggested a different cause. Albeit neutrophils were indeed less deformable and smaller 15 min post fMLP treatment, the subsequent time-course showed reversal to more deformed and larger cells (Fig. 3A, B; Fig. S7). So, the likely reason for the discrepancy of previous reports could lie in the different time points of measurement after activation.

**Fig. 3.**
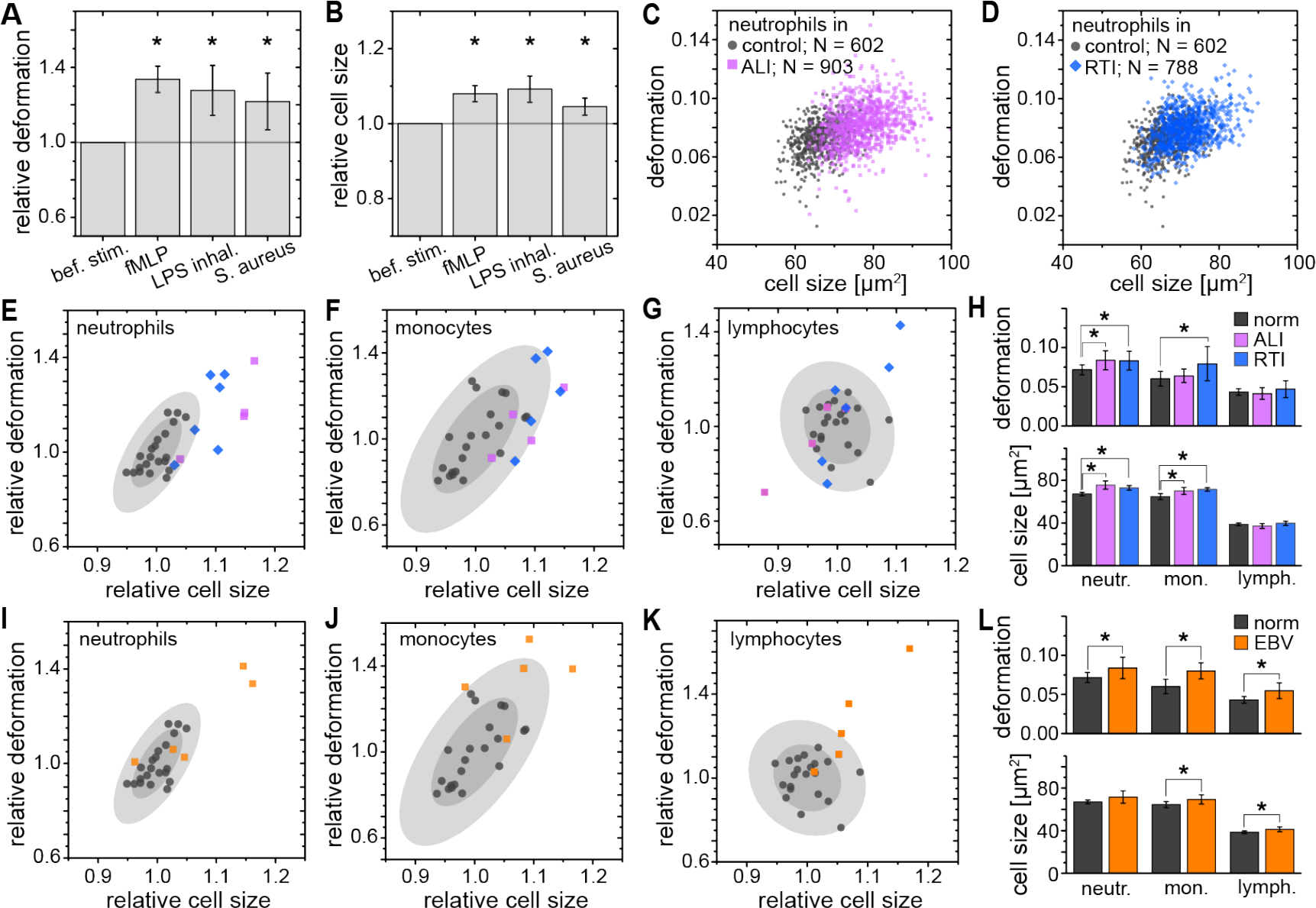
MORE phenotyping identifies leukocyte activation and infections *in vitro* and *in vivo*. Relative change (mean ± SD) in **A**, deformation and **B**, size of neutrophils in whole blood after fMLP (*n* = 5) and *S. aureus* (*n* = 4) stimulation *in vitro*, and LPS inhalation (*n* = 2) *in vivo*. Exemplary scatter plots of size vs. deformation of neutrophils in blood of a patient with **C**, ALI (magenta) and **D**, RTI (blue) compared to controls (black). Medians of size and deformation of **E**, **I**, neutrophils, **F**,**J**, monocytes,and **G**, **K**, lymphocytes in blood samples of patients with E, F, G, ALI (*n* = 4) and RTI (*n* = 6), and I, J, K,EBV infection (*n* = 5; orange) relative to the norm (black, *n* = 21 as in Fig. 1E with 68 % and 95 % confidence ellipses). **H**, **I**, Mean and SD of these results, * *p* < 0.05.

We found the same softening response in an experimental medicine trial, where lipopolysaccharide (LPS; from *E. coli*) inhalation in healthy human volunteers likewise induced a transient increase in size and greater deformation of the neutrophils (Fig. 3A, B;Fig. S7). Also, infecting whole blood *in vitro* with *Staphylococcus aureus (S. aureus)*, a Gram-positive bacterium and one of the major pathogens responsible for life-threatening infections world-wide, resulted in larger and more deformed neutrophils, measured between 30 – 60 min after blood stimulation (Fig. 3A, B; Fig. S8).

Congruently, blood taken from patients with an acute lung injury (ALI) of most likely bacterial origin had larger and more deformed neutrophils compared to healthy controls (Fig. 3C, E, H). The same neutrophil response was found in blood samples from patients hospitalized with viral respiratory tract infections (RTI; Fig. 3D, E, H). Also monocytes responded by a size increase in both RTI and ALI patients and after *in vitro* stimulation with *S. aureus*, but only in viral RTI showed a significantly increased deformation, while blood lymphocytes did not show any consistent response (Fig. 3F-H; Fig. S8). The lymphocyte response changed when analyzing blood of patients with acute Epstein-Barr-virus (EBV) infection, which is known to also stimulate the lymphatic system, where both monocytes and lymphocytes showed an increase in cell size and deformation, while neutrophils showed less of a response (Fig. 3I-L). These results suggest that MORE blood analysis might be sufficiently sensitive to distinguish bacterial from viral infections, and potentially other inflammatory diseases, by the differential response of the selective blood leukocyte populations. This possibility will be followed up in future specific trials. Importantly, MORE blood analysis is of special interest for blood tests in neonatology with patients at high risk of infections but only minute amounts of blood available for diagnostics, or to characterize neutrophils in neutropenic patients, as it merely requires longer data acquisition periods.

### MORE analysis of malignant transformed blood cells

Blood cancers, or leukemias, affecting both myeloid and lymphoid cell lineages, are a further large area, where MORE analysis could potentially contribute fundamental insight, aid diagnosis, and improve therapy monitoring. While solid cancer cell mechanics has been a focus of cell mechanics research and extensively documented (27–29), the mechanical properties of blood cancers are comparatively understudied. The available mechanics research on leukemic cells has been undertaken either on cell lines or fully purified cells (1, 7–10, 12, 13, 17, 30) but so far not in whole blood. MORE analysis of the blood of patients with acute myeloid (AML) and lymphatic leukemias (ALL) revealed the new presence of atypical cell populations — the characteristic immature blasts not normally present in healthy donors (Fig. 4A-C). Cell populations gated for AML revealed less deformed cells but at about the same size compared to healthy and fully differentiated myeloid cells (Fig. 4D, Fig. S9), in line with previous results (7, 9, 10, 12, 13).

**Fig. 4.**
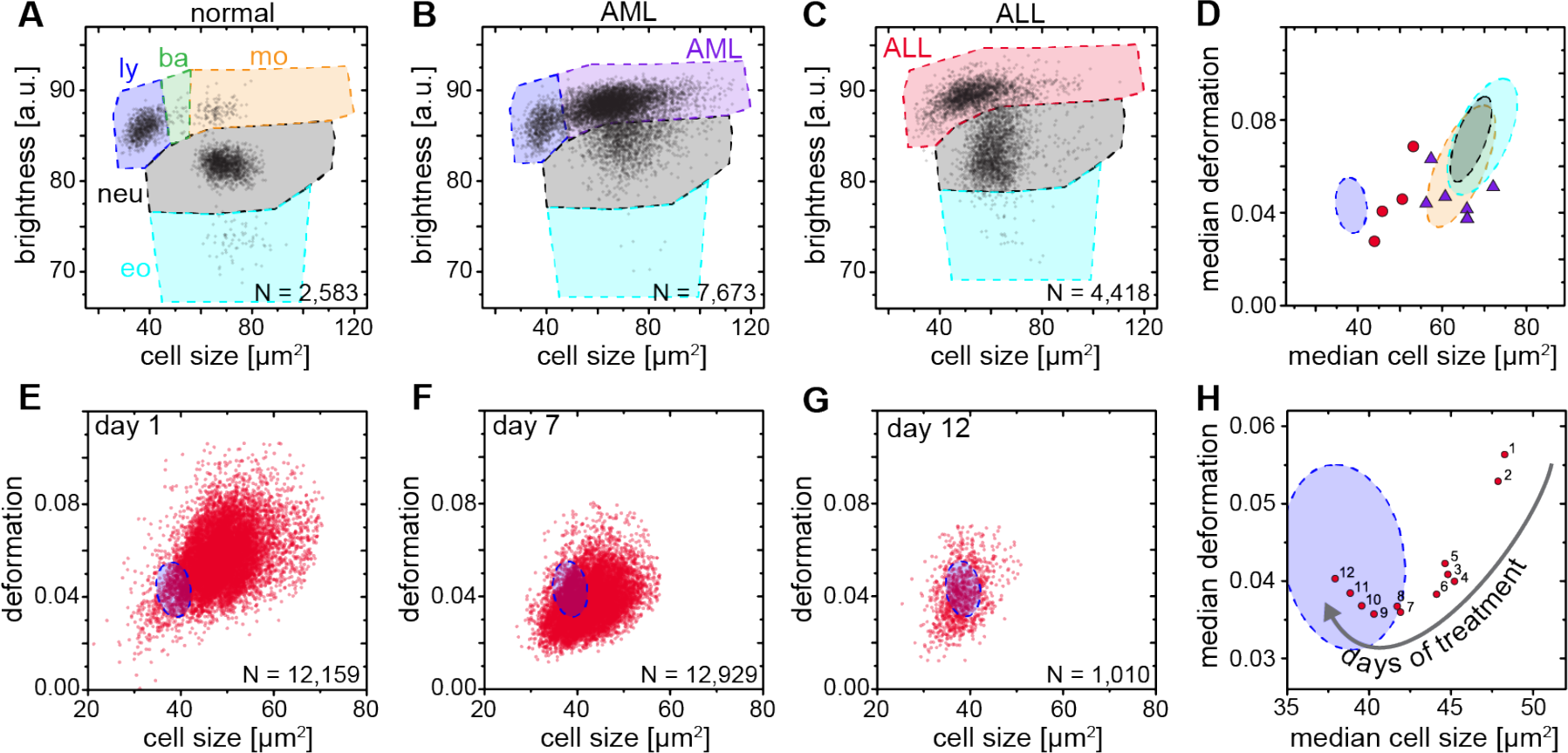
MORE phenotyping detects and distinguishes leukemia subtypes and monitors treatment effects. **A**, Normal brightness vs. size scatter plot of a healthy donor with the gates (shaded areas) used to identify lymphocytes (ly), basophils (ba), monocytes (mo), neutrophils (neu) and eosinophils (eo). **B**, Exemplary brightness vs. size scatter plot in AML; blast cells were found in (ba) and (mo) gates. **C**, Exemplary brightness vs. size scatter plot in ALL; blast cells were found in (ly), (ba), and (mo) gates. **D**, Medians of deformation and size for the respective gates in blood samples of ALL (red circles, *n* = 4) and AML (purple triangles, *n* = 7) patients. Shaded areas in D-H (color as in A) represent 95% confidence ellipses of the respective cell type norm (*n* = 21, as in Fig. 1E). Scatter plots of ALL blast deformation and size at **E**, one; **F**, seven, and **G**, twelve days post therapy start. **H**, Median deformation and size of ALL cells during 12 days of treatment (red dots).

ALL blast cells were larger in size compared to mature lymphocytes, but did not show any consistent trend in deformation (Fig. 4D; Fig. S9). Since larger cells of identical stiffness should deform more in RT-DC (20, 21, 31), these findings together imply that mature lymphocytes, ALL blasts, mature myeloid cells, and AML blasts have decreasing levels of deformability, consistent with the composite findings of previous reports (1, 7–10, 12, 13, 17). This is quite different to the general trend in solid tumors, where cancer cells are found to be more deformable than their healthy counterparts (27–29). Sensibly, the differential stiffness of AML and ALL blasts, and its potential further increase with chemotherapy, has been implicated in the occurrence of leukostasis (8, 17, 32). MORE analysis might not only permit screening for novel therapeutic targets to soften cells (19, 33, 34), but also assessing the risk of leukostasis directly in each patient. In addition, by following the ALL blast population in a patient over 12 days of methylcortisone treatment we could monitor the return to the normal morpho-rheological fingerprint of blood as mature lymphocytes successfully replaced the lymphoblasts (Fig. 4E-H). Thus, MORE blood analysis can be used to monitor morpho-rheological effects of chemotherapy in a quantitative manner. This last finding also touches upon the study of hematopoietic differentiation of cells in the bone marrow, which is an obvious further potential area of application of this approach.

## Discussion and Conclusion

MORE phenotyping allows individual blood cell mechanics to be studied in a range of human diseases and takes cell mechanical phenotyping to an entirely new level. While established techniques such as micropipette aspiration (1, 9–11), indentation by cell pokers and atomic force microscopes (6–8), or optical trapping (12–14) have provided important proof-of-concept insight over the last decades, the recent advent of microfluidic techniques approaching the throughput of conventional flow cytometers (19, 21, 30, 35, 36) has finally brought mechanical phenotyping close to real-world applications (29, 37). Amongst the latter techniques, RT-DC stands out because it can continuously monitor an unlimited number of cells, which enables the direct sensitive assessment of the state of all major blood cell types directly in whole blood. A volume as small as 10 μl can be analyzed cell-by-cell, with only minimal dilution and no labeling, enrichment or separation, which could otherwise cause inadvertent activation of blood cells. The conventional blood count is extended by information about characteristic, and diagnostic, morpho-rheological changes of the major cell types. Cell mechanics and morphology are inherent and sensitive markers intimately linked to functional changes associated with the cytoskeleton (38–42) and other intracellular shape-determining and load-bearing entities (43, 44). Thus, label-free, disease-specific MORE blood signatures are a novel resource for generating hypotheses about the underlying molecular mechanisms. The availability of such parameters in real-time, easily combined with conventional fluorescence detection, are the necessary prerequisite for future sorting of morpho-rheologically distinct subpopulations, which then provides a novel opportunity for further molecular biological analysis. Of course, at present, MORE phenotyping provides a sensitive, but not a very specific marker. For example, neutrophil softening could be a signature of different underlying pathological changes. In the future, fuller exploration of the large combinatorial space afforded by the multi-parametric response of the various blood cells, exploiting many additional MORE parameters in conjunction with machine learning, and inclusion of conventional fluorescence-based marker analysis will further increase the specificity of this approach. Apart from now enabling realistic blood cell research *ex vivo* close to physiological conditions, delivering for example previously unavailable information about leukocyte activation kinetics, and after further in depth studies of the phenomena reported here, MORE phenotyping could have a tangible impact on diagnosis, prognosis, and monitoring of treatment success of many hematological diseases as well as inflammatory, infectious, and metabolic disorders. Beyond blood analysis, MORE phenotyping has the potential to become a standard approach in flow cytometry with many applications in biology, biophysics, biotechnology and medicine.

## Materials and Methods

### Real-time deformability cytometry

Real-time deformability cytometry (RT-DC) was carried out as described previously (21). For RT-DC measurements cells were suspended in a viscosity-adjusted measurement buffer (MB) based on 1x phosphate buffered saline (PBS) containing methylcellulose. The viscosity was adjusted to 15 mPa s at room (and measurement) temperature, determined using a falling ball viscometer (Haake, Thermo Scientific). Cells in the MB were taken up into a 1 ml syringe, placed on a syringe pump (neMESYS, Cetoni GmbH) and connected via tubing to the sample inlet of the microfluidic chip with a square measurement channel cross section of 20 × 20 μm^2^. The microfluidic chip was made from cured polydimethylsiloxane bonded to a thickness #2 cover glass. Another syringe containing MB without cells was connected to the sheath flow inlet of the chip. Measurements were carried out at a total flow rate of 0.12 μl/s with a sample flow rate of 0.03 μl/s and a sheath flow rate of 0.09 μl/s unless stated otherwise. Different gating settings for cell dimensions could be employed during the measurement (Fig. S1). Images of the cells in the channel were acquired in a region of interest of 250 × 80 pixels at a frame rate of 2,000 fps. Real-time analysis of the images was performed during the measurement and the parameters necessary for MORE analysis were stored for all detected cells.

### Data processing in MORE analysis

The raw data obtained from RT-DC measurements consisted of the following information of every detected cell: a bright field image of the cell, the contour of the cell, its deformation value, and the cell size as the cross-sectional area of the cell in the image (Fig. S1). The deformation was calculated from the convex hull contour of the cell — a processed contour, where all points contributing to concave curvature were removed:

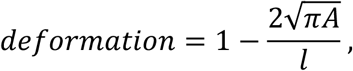

where *A* is the area enclosed by the convex hull contour and *l* is the length of the convex hull contour. Therefore, deformation is the deviation from a perfectly circular cell image. It describes the change of the cell’s shape by the hydrodynamic forces in the measurement channel but may also contain pre-existing shape deviations from a sphere. Image brightness analysis was carried out using the contour information and the image of the cell. The mean brightness of the cell was determined from all pixel values within the cell’s contour (Fig. 1D). With this information the distinction of leukocyte subpopulations was possible in the space spanned by cell size and mean brightness (Fig. 1D and Fig. S2). It is worth noting that the absolute value of the resulting brightness was influenced by several experimental conditions such as focus of the image and the thickness of the microfluidic chip. However, this did not affect the quality of the distinction of cells by their brightness. Special care had to be taken when comparing the brightness of different purified leukocyte subpopulations of similar size (like neutrophils, eosinophils, and monocytes). In order to achieve a situation similar to the whole blood measurement we used the same microfluidic chip repeatedly after thorough flushing. All brightness values reported were normalized to 100 by the background brightness of the channel. Apart from the initial brightness distinction, in a second step, the root mean square of pixel brightness values was calculated in an area of 9 × 5 pixels (9 in the flow direction, 5 perpendicular to the flow direction) around the geometrical center of the cell. This information was used to distinguish the relevant leukocyte subpopulations from eventual erythrocyte doublets present (Fig. 1D). To ensure best validity of the deformation measure based on the area within the cell’s contour and the length of the contour, only cells without prominent protrusions were considered for comparisons based on deformation. A reliable criterion to select those cells was found by comparing the area within the originally detected cell contour and within the convex hull contour. For erythrocytes, the difference of these two areas was limited to 15 %. For leukocytes, a suitable limit was found at 5 %. For the identification of malaria-infected erythrocytes we used a semi-automated procedure designed to obtain only clearly positive results and to avoid false negatives. The defining property of infected cells was the presence of bright spots within the cells. In a first step, all pixel values outside the cell’s contour were set to 0. In a twice-repeated procedure, the image of the erythrocyte was further reduced by setting all pixel values of the contour pixels to 0 and finding the new contour. This measure was used to eliminate possible bright spots due to fringes at the border of the cell. From this image, the brightness of every pixel of the remaining cell was calculated by taking the mean of the pixel itself and its 8 nearest neighbors. The user was then able to set the minimal threshold for this brightness in order to identify a cell as potentially infected. Since higher pixel values are frequently obtained at the rear of the cell (in flow direction) only bright spots within 70 % of the cell’s length counted from the front of the cell were considered. As a last criterion, the calculated brightness was compared to the brightness of the cell directly surrounding the bright spot in order to eliminate cases of generally bright cells. For this a mean brightness value was formed from all pixels located within the two rectangular areas spanned from [*k*−3,*l*−1] to [*k*−2,*l*+1] as well as [*k*+2,*l*−1] to [*k*+3,*l*+1], where *k* is the pixel position of the bright spot in the flow direction and *l* is the pixel position of the bright spot orthogonal to the flow direction. All scripts for MORE analysis were written in Matlab and Python using OpenCV.

### Whole blood measurements

All studies complied with the Declaration of Helsinki and involved written informed consent from all participants or their legal guardians. Ethics for experiments with human blood were approved by the ethics committee of the Technische Universität Dresden (EK89032013, EK458102015), and for human blood and LPS inhalation in healthy volunteers by the East of England, Cambridge Central ethics committee (Study No. 06/Q0108/281 and ClinicalTrialReference NTC02551614). Study participants were enrolled according to good clinical practice and recruited at the University Medical Centre Carl Gustav Carus Dresden, Germany, the Biotechnology Center, Technische Universität Dresden, Germany, or Cambridge University Hospitals, Cambridge, UK. Human blood and serum used to culture the malaria parasites was obtained from the Glasgow and West of Scotland Blood Transfusion Service; the provision of the material was approved by the Scottish National Blood Transfusion Service Committee For The Governance Of Blood And Tissue Samples For Non-Therapeutic Use. Venous blood was drawn from donors with a 20-gauge multifly needle into a sodium citrate S-monovette (Sarstedt) by vacuum aspiration. In case of blood volumes above 9 ml, blood was manually drawn via a 19-gauge multifly needle into a 40 ml syringe and transferred to 50 ml Falcon polypropylene tubes (BD) containing 4 ml 3.8% sodium citrate (Martindale Pharmaceuticals). For whole-blood RT-DC measurements, 50 μl of anti-coagulated blood were diluted in 950 μl MB and mixed gently by manual rotation of the sample tube. Measurements were typically carried out within 2 h past blood donation unless stated otherwise. Two different gating settings were employed in the measurement software for erythrocyte and leukocyte acquisition, respectively (Fig. S1A). For erythrocytes, gates were essentially open allowing cell dimensions in flow direction from 0 μm to 30 μm. The leukocyte gate was set to a size of 5 – 16 μm in flow direction and > 5 μm perpendicular to it. This setting allowed filtering out single erythrocytes and almost all erythrocyte multiples. The leukocyte populations remained unaltered as confirmed in experiments with purified leukocytes at open gate settings. Using the leukocyte gate, the majority of thrombocytes was also ignored as they possess typical diameters of 2 – 3 μm. A small fraction of very large thrombocytes and microerythrocytes were still found within this gate as seen in Fig. 1C and D. Mechanical analysis of these events constitutes an interesting challenge in that they can be detected and counted, but at present not tested for activation via their deformation given their very small size compared to the channel size, which was chosen to accommodate all cells found in blood. Measurements in the leukocyte gate were carried out over a timespan of 15 min, followed by a separate measurement in the erythrocyte gate for a few seconds until data of 5,000 – 10,000 cells were acquired. Measurements for establishing the normal MORE blood phenotype in healthy human volunteers (Fig. 1E), and all measurements directly compared to this norm, e. g., blood samples derived from patients, were carried out at a temperature of 30 °C. The remaining measurements — fMLP stimulation, LPS stimulation, purified leukocyte subpopulations, malaria infection, and erythrocyte palmitoylation — were carried out at a temperature of 23 °C. The viscosity of the MB was always adjusted to 15 mPa s at the different temperatures to keep the acting hydrodynamic stress and, thus, the resulting deformation regimes the same. An MB with the viscosity of 25 mPa s (to slow blood cell sedimentation in the tubing) was used in experiments for comparing the relative cell count results of leukocyte subpopulation by MORE analysis and conventional blood count (Fig. 1F; Table S1). Here, the total flow rate was 0.06 μl/s (sample flow 0.015 μl/s, sheath flow 0.045 μl/s) and images were acquired at 4,000 fps.

### Leukocyte purification and identification

Leukocyte subpopulations were purified by negative and/or positive magnetic-activated cell sorting (MACS) following the instructions provided by the manufacturer. Reagents for cell isolation with magnetic beads purchased from Miltenyi Biotec were MACSxpress Neutrophil Isolation Kit human (130-104-434), Monocyte Isolation Kit human (130-091-153), Basophil Isolation Kit II human (130-092-662), Pan T Cell Isolation Kit human (130-096-535) and CD3 MicroBeads (130-050-101), as well as Pan B Cell Isolation Kit human (130-101-638) and CD19 MicroBeads (130-050-301). EasySep Human Eosinophil Enrichment Kit (19256) was obtained from StemCell Technologies. The purity of the derived cell isolates was controlled twice by staining with 7-Color-Immunophenotyping Kit (Miltenyi Biotec, 5140627058), as well as additional single staining of each cell subset for fluorescence-activated cell sorting (FACS). Individual cell type staining antibodies from BioLegend were used for granulocytes (target: CD66ACE, staining: PE, order no.: 342304), eosinophils (Siglec-8, APC, 347105), B lymphocytes (CD19, FITC, 302205), NK cells (CD56, PE, 318305), T helper cells (CD4, PE-Cy7, 300511), T lymphocytes (CD3, APC, 300411), cytotoxic T cells (CD8, PacificBlue, 301026), monocytes (CD14, FITC, 325603), as well as eosinophils, basophils, mast cells, and mononuclear phagocytes (CD193, PE, 310705). For RT-DC measurements, purified cells were pelleted by centrifugation (200 g, 5 min) and re-suspended in MB at concentrations of about 5 · 10^6^ cells/ml by repeated, gentle shaking.

### In vitro malaria infection

*Plasmodium falciparum* (*P. falciparum*) cultures were grown accordingly to standard protocols (45). Two *P. falciparum* cultures (HB3 clone), were grown independently for 3 weeks, treated with Plasmion(46) to enrich for the schizont stages, and then allowed to reinvade fresh red blood cells in a shaking incubator for 3 h. The cultures were then treated with sorbitol (47), to remove all schizonts that had not reached full maturity; only ring stage parasites survive sorbitol treatment. The highly synchronised culture used for the RT-DC measurements therefore consisted of erythrocytes into which parasites had invaded within a 3 h window. Samples were removed at 4, 12, 16, 20, 24, 36, 42 and 46 hours post invasion for the RT-DC measurements. At the time of each measurement a thin blood smear was taken and stained with Giemsa’s stain to assess the parasitemia and the stage of the parasites (Fig. S6A). A control sample of the same blood without the parasites underwent the identical treatment as the infected samples. For RT-DC measurements, at each time point, 10 μl of the blood culture were diluted in 990 μl of the MB to a final concentration of 2.5 · 10^5^ cells/ μl. The total flow rate through the channel was 0.04 μl/s for all malaria infection experiments (sample flow rate 0.01 μl/s, sheath flow rate 0.03 μl/s). For experiments on growth and invasion depending on erythrocyte palmitoylation status, blood, treated as described in the palmitoylation section below, was shipped from Germany to Scotland in PBS buffer containing 15 mM glucose, 5 mM sodium pyruvate, 5 μM Coenzyme A, 5 mM MgCl_2_, 5 mM KCl, 130 mM NaCl. Parasites were synchronized by collecting *P. falciparum* mature stages (trophozoites and schizonts) from *P. falciparum* clone HB3 using MACS columns(48). The trophozoite and schizont enriched cultures were mixed with erythrocytes to achieve a starting parasitemia of 0.5 – 1.0 %. Each erythrocyte type was set up in a separate culture flask at 3 ml volume and 5 % hematocrit. The parasites were incubated in a shaking incubator at 37°C under standard culture conditions of gas and medium. Parasitemia was monitored on day 2 (post invasion) and day 4 (second round of invasion). For all experimental conditions, a minimum of 500 RBCs were counted. Experiments were repeated on 3 different days with erythrocytes of 3 different donors yielding the same results.

### Palmitoylation of erythrocytes

Red blood cells were pelleted by blood centrifugation (800 g, 5 min), plasma was removed, and the RBCs were pretreated with one volume of 1 % fatty acid-free bovine serum albumin (BSA) in PBS-glucose (10 mM phosphate, 140 mM NaCl, 5 mM KCl, 0.5 mM EDTA, 5 mM glucose, pH7.4) at 37° C for 15 min, in order to lower the endogenous content of free fatty acids in their membrane pools, and washed three times with PBS-glucose. Cells were re-suspended in 3 volumes of incubation buffer, containing 40 mM imidazole, 90 mM NaCl, 5 mM KCl, 5 mM MgC12, 15 mM D-glucose, 0.5 mM EGTA, 30 mM sucrose, 5 mM sodium pyruvate, 5 mM Coenzyme A, 50 mg PMSF/ml and 200 U penicillin/streptomycin (320 mOsm, pH 7.6). For inhibition of palmitoylation, 100 μM final concentration of 2-bromopalmitate (2BP) was used. 100 μM palmitic acid (PA) was added as a control. The RBCs were incubated in a humidified incubator with 5 % CO_2_ for 24 h at 37 °C. Prior to measurement, RBCs were pelleted, re-suspended in 1 % BSA, incubated for 15 min at 37 °C and washed two times with PBS-glucose. Glucose, sucrose, 2-bromopalmitate, palmitic acid, fatty acid free BSA, Coenzyme A, and PMSF were purchased from Sigma-Aldrich; Penicilin/streptomycin and sodium pyruvate from Gibco. RT-DC measurements were carried out at a room temperature of 23°C and with a total flow rate of 0.032 μl/s (sample flow 0.008 μl/s, sheath flow 0.024 μl/s) after adding 10 μl of the RBC suspension to 990 μl of MB. Experiments were carried out on 2 different days with erythrocytes of 4 different donors.

### fMLP-induced neutrophil activation

For *in vitro* fMLP stimulation, whole blood was stimulated with 100 nM N-Formylmethionyl-leucyl-phenylalanine (fMLP; Sigma-Aldrich, 47729, 10 mg-F). Separate samples were analyzed in time intervals of 0 – 15 min, 15 – 30 min, 30 – 45 min, and 45 – 60 min after activation. During incubation all samples were stored in 2 ml Eppendorf tubes at 37°C at 450 rpm in a ThermoMicer C (Eppendorf). All experiments were performed within 2 hours maximum after blood drawing. Experiments were repeated with blood samples of 5 different donors on 5 different days. Due to experimental feasibility PBS controls of these donors were measured before fMLP stimulation and after the 60 min fMLP sample. Additionally, three control samples of different donors were treated similarly adding 10 μl 1 × PBS instead of fMLP and were analyzed in time intervals of 0 – 15 min, 15 – 30 min, 30 –45 min, and 45 – 60 min after bleeding to exclude kinetic effects due to blood alteration with storage.

### In vitro Staphylococcus aureus infection

Whole blood stimulation was performed with *Staphylococcus aureus* Newmann strain (*S. aureus*; ATCC25904). For reproducible repetitive testing with competent bacterial strains cryo-aliquots of *S. aureus* were prepared as follows. Bacterial cells were pre-cultured to the log phase for synchronization of growth in BHI broth (Bacto Brain Heart Infusion, Becton Dickinson) at 37°C and transferred to a second culture. Aiming at a high bacterial virulence factor expression, the cells were grown to an early stationary phase in a 96-well-plate (100 μl, OD_600nm_ 0.1837, Infinite 200 reader, TECAN), pelleted by centrifugation (2671 g for 5 min at 4 °C), washed two times in PBS and re-suspended in cell-freezing media (Iscove Basal Medium, Biochrom) with 40 % endotoxin-free FBS (FBS Superior, Biochrom) at a final concentration of 2.54⋅10^9^ CFU/ml. Aliquots were immediately frozen at –80 °C and only thawed once for a single experiment. Blood stimulation and measurement were carried out at 30 °C temperature for 15 min with one multiplicity of infection (MOI) in 1:20 RT-DC measurement buffer. MOI (0.9 - 1.09) was controlled retrospectively by granulocyte count and 5 % sheep blood agar culture (Columbia agar, bioMérieux) at 37 °C and bacterial colony counting on the following day. PBS blood controls were conducted before and after *S. aureus* blood stimulation. The experiment was repeated with blood of 4 different donors on 4 different days. All experiments were performed within 2 h after blood drawing.

### LPS inhalation

*E. coli* lipopolysaccharide (LPS) 50 μg (GSK) was administered to healthy, never-smoker volunteers via a specialized dosimeter (MB3 Markos Mefar) 90 minutes prior to injection of autologous ^99m^Technetium-Hexamethylpropleneamine-oxime labeled neutrophils. Temperature, forced expiratory volume in 1 second, forced ventilator capacity and triplicate blood pressures were recorded prior to, and at 30 min intervals post LPS administration. RT-DC measurements were obtained at baseline, 90, 135, 210, 330, and 450 min post LPS.

### Respiratory tract infections (RTI) and acute lung injury (ALI)

Patient inclusion criteria for RTI: Patients with clinical signs of lower RTI, a core temperature > 38.5°C and the need for supplemental oxygen were recruited on the day of hospitalization. Only patients without treatment prior to hospitalization were included. None of the included patients received antibiotic treatment for reconstitution. Patient inclusion criteria for ALI: Patients diagnosed with ALI according to the criteria of the North American European Consensus Conference (NAECC) (49) and without underlying diseases prior to ALI were included. All blood samples were analyzed within 30 min of venipuncture. Size and deformation of blood leukocytes was characterized for all blood cells in which the area within the original cell contour differed less than 5 % from the area within the convex hull contour.

### Acute myeloid/lymphatic leukemias

Samples from patients diagnosed with ALL or AML based on cytogenetic, molecular-genetic and morphological criteria according to WHO classification from 2008 (50) were assessed by MORE blood analysis on the day of diagnosis. In order to evaluate mechanical properties of AML and ALL blast cells in whole blood, several brightness and size gates had to be combined as shown in Fig. 4A-C. The AML gate spanned the regions normally used for basophils and monocytes. The ALL gate spanned the regions used for lymphocytes, basophils and monocytes. In all AML cases, blasts made up > 80 % of all leukocytes, and up to 99 % of events in the AML gate. In all ALL cases, blasts made up > 60 % of leukocytes, and up to 85 % of events in the ALL gate. The blast cell fraction was obtained from the standard differential blood count, by comparing the number of blast cells with the number of normal cells that would also populate the respective blasts gate in MORE analysis.

### Statistics

Throughout, the number of cells in a single measurement is denoted as *N*, while the number of repeated experiments is denoted as *n*. For comparison of different donors or treatment conditions the median of deformation and cell size of a specific cell population was used. In order to evaluate effects of a disease we calculated a 2D confidence ellipse at 68.3 % (or 1 sigma) as well as 95.5 % (or 2 sigma) for the control group/norm norm of healthy human blood donors in the space of cell size and deformation. The confidence ellipse was calculated from the covariance matrix of the data and the calculation was carried out with OriginPro 2015 (Originlab). Statistically significant differences between two sets of experiments were checked to the significance level of *p* < 0.05 by comparing the groups of individual median values of an experiment using a Kruskal-Wallis one-way ANOVA as implemented in OriginPro 2015 (Originlab). In erythrocyte MORE analysis in malaria infection and palmitoylation, statistically significant differences were checked using linear mixed models (51) in combination with a likelihood ratio test to obtain significance levels for the comparison of the complete populations. One, two, or three asterisks were awarded for significance levels *p* < 0.05, *p* < 0.01 and *p* < 0.001, respectively. In manual counts of malaria infection in RBCs, statistical analyses were performed using a χ^2^ test with Bonferroni correction (adjusted statistical significance for *p* < 0.0125)to compare the numbers of infected and non-infected erythrocytes between erythrocyte samples, except where number of parasite infected cells was zero, in which case Fisher’s exact test was used. The standard deviation for the parasitemia was calculated assuming a binomial random variable as 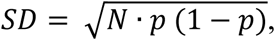 where *N* is the number of cells counted and *p* is the fraction of infected cells.

### Code availability

RT-DC measurement software is commercially available. MORE analysis software is available as an open source application at https://github.com/ZELLMECHANIK-DRESDEN/ShapeOut/releases.

### Data availability

The data supporting the findings of this study are available from the corresponding author upon reasonable request.

## Acknowledgements

The authors would like to thank Björn Lange, Michael Mögel, Beate Eger, Isabel Deinert, Tamara Schön and the whole team for their contributions to patient recruitment, Claudia Krug for cell isolation, Uta Falke and Isabel Richter for technical help, Thomas Krüger for help with drawing blood, Ramona Hecker for technical advice, Mike Blatt for the loan of a microscope, Elizabeth Peat for assistance with malaria parasite culture materials, Salvatore Girardo of the BIOTEC/CRTD Microstructure Facility (in part funded by the European Fund for Regional Development – EFRE) for help with preparation of PDMS chips, and Stephan Grill for critical reading of the manuscript.

Financial support from the Alexander von Humboldt Stiftung (Alexander von Humboldt Professorship to J.G.), a CRTD Seed Grant (A.J. and J.G.), the FP7-funded ITN “LAPASO” (L.C., L.R.-C., E.R., G.G., P.M., B.H.-N., and J. G.),TUD Support the Best (R.B and J.G.), TG70 (O.O. and J.G.), the ERC Starting Grant “LightTouch” (J.G.), the DFG TRR83 grant TP18 (M.G. and Ü.C.), the BMBF grant to the German Center for Diabetes Research (DZD e.V.) (M.G. and Ü.C.), the DFG SFB655 grant ‘From cells to tissues’ (subproject B2 to M.K. and M.B.), non-commercial grants from ‘Tour der Hoffnung’ (J.S.) and ‘Sonnenstrahl e.V. Dresden’ (M.S.), and GlaxoSmithKline (N.Tr. and the LPS inhalation study), and the NIHR Cambridge Biomedical Research Centre (E.R.C) is gratefully acknowledged.

## Author contributions

N.To. and C.H. conceived the project and designed and carried out most experiments, developed analysis methods, interpreted results and co-wrote the manuscript. O.O. and P.R. guided and performed technical developments of the RT-DC and helped with the experiments and analysis. A.J., M.K., J.S., L.M. assisted cell isolation experiments and performed the leukemia experiments. M.H. programmed analysis methods. L.C. and L.R.C. conceived, performed and coordinated the malaria experiments. M.G. and Ü.C. conceived, performed and coordinated the palmitoylation experiments. E.R., G.G., P.M. contributed to the experiments on bacterial neutrophil activation. E.R.C. and N.Tr. conceived the experiments on LPS inhalation. E.R.C., B.H.N., M.S., M.B. and R.B. provided intellectual contributions and contributed towards the manuscript writing. J.G. conceived the project, supervised experimental designs, interpreted results, and co-wrote the manuscript.

## Declaration of potential conflict of interest

C.H., O.O and P.R. own shares of, and are part- or full-time employed at, Zellmechanik Dresden GmbH, a company selling devices based on real-time deformability cytometry.

